# Natural selection drives population divergence for local adaptation in a wheat pathogen

**DOI:** 10.1101/805127

**Authors:** Danilo Pereira, Daniel Croll, Patrick C. Brunner, Bruce A. McDonald

## Abstract

Evolution favors the emergence of locally-adapted optimum phenotypes that are likely to differ across a wide array of environmental conditions. The emergence of favorable adaptive characteristics is accelerated in agricultural pathogens due to the unique properties of agro-ecosystems. We performed a Q_ST_ - F_ST_ comparison using 164 strains of *Parastagonospora nodorum* sampled from eight global field populations to disentangle the predominant evolutionary forces driving population divergence in a wheat pathogen. We used digital image analysis to obtain quantitative measurements of growth rate and melanization at different temperatures and under different fungicide concentrations in a common garden experiment. F_ST_ measures were based on complete genome sequences obtained for all 164 isolates. Our analyses indicated that all measured traits were under selection. Growth rates at 18°C and 24°C were under stabilizing selection (Q_ST_ < F_ST_), while diversifying selection (Q_ST_ > F_ST_) was the predominant evolutionary force affecting growth under fungicide and high temperature stress. Stabilizing selection (Q_ST_ < F_ST_) was the predominant force affecting melanization across the different environments. Melanin production increased at 30°C but was negatively correlated with higher growth rates, consistent with a trade-off under heat stress. Our results demonstrate that global populations of *P. nodorum* possess significant evolutionary potential to adapt to changing local conditions, including warmer temperatures and applications of fungicides.

## 1. INTRODUCTION

Evolution by means of natural selection operates on individual phenotypes and is enabled by the diversity found in genes encoding quantitative traits within populations. Across a wide array of environmental conditions, evolution towards a local optimum phenotype results from the interplay among evolutionary forces such as natural selection and gene flow and stochastic events like founder events and population extinctions (Merilä and Crnokrak, 2001; Leinonen et al., 2008). Evolutionary processes affecting local adaptation of pathogen populations may act differently in agro-ecosystems compared to natural environments. For example, the high planting densities found in agricultural fields allow more efficient pathogen transmission and the genetic uniformity of agricultural hosts enable the development of large pathogen populations while imposing strong directional selection that accelerates the emergence of host specialization (Stukenbrock and McDonald, 2008; McDonald and Stukenbrock, 2016; Corredor-Moreno and Saunders, 2019). While particular agro-ecosystems (e.g. the one used for wheat production) tend to be highly similar on a global spatial scale, local pathogen populations can encounter significant differences in the deployment of resistance genes, pesticide exposure and annual fluctuations in temperature over a growing season (Laine, 2008; Stukenbrock and McDonald, 2008; Elderd and Reilly, 2014). Hence, we expect that even globally distributed pathogens can evolve different traits in different local populations according to the predominating local evolutionary forces.

A better understanding of which evolutionary forces are driving local adaptation for a particular trait can be achieved using Q_ST_ - F_ST_ comparisons (Zhan et al., 2005; Leinonen et al., 2008; Leinonen et al., 2013; Stefansson et al., 2014; Yang et al., 2016). Q_ST_ is an index of population differentiation based on the distribution of variation for a quantitative trait (Spitze, 1993). F_ST_ measures the degree of population divergence based on neutral genetic markers. Natural selection is inferred when population differentiation for quantitative traits is significantly different from that for neutral markers (Q_ST_ ≠ F_ST_). Specifically, directional selection is inferred when the Q_ST_ is higher than the F_ST_, and stabilizing selection is inferred when Q_ST_ is lower than the F_ST_ (Leinonen et al., 2013). When no differences are found between the two indexes, the inference is that the trait is neutral or that it is not possible to distinguish between the effects of genetic drift and natural selection in the populations being examined. Previous Q_ST_ - F_ST_ comparisons for plant pathogenic fungi were conducted using global field populations of the wheat pathogen *Zymoseptoria tritici* and the barley pathogen *Rhynchosporium commune.* In both cases, natural selection was inferred to be the main driver of local adaptation (Zhan et al., 2005; Stefansson et al., 2014), but directional selection predominated in *Z. tritici,* while stabilizing selection was more important in *R. commune.* These differences highlight the necessity to consider the adaptive dynamics of each trait in a species-specific manner.

The fungal pathogen *Parastagonospora nodorum* causes stagonospora nodorum blotch (SNB), a major wheat disease found around the world (Quaedvlieg et al., 2013; Savary et al., 2019). Field populations of *P. nodorum* are reported to have low genetic differentiation among continents, elevated population size, high genetic diversity and exhibit frequent sexual recombination (Stukenbrock et al., 2006; Oliver et al., 2012). SNB control measures include fungicide applications and the deployment of wheat varieties that lack toxin sensitivity genes (Oliver et al., 2012; Ficke et al., 2017). Fungicides belonging to the sterol demethylation inhibitors group (DMIs) are commonly used in both agriculture and human medicine (Price et al., 2015). DMI-resistant strains of *P. nodorum* harboring point mutations in the gene encoding the targeted protein (CYP51) have been previously reported (Pereira et al., 2017). While the severity of SNB is influenced by environmental factors (e.g. SNB is most damaging in warm and moist conditions (Shaw et al., 2008; Zearfoss et al., 2011), the costs of managing SNB were estimated to be AUD$108 m per year in Australia alone (Murray and Brennan, 2009). A better understanding of the evolutionary processes affecting local adaptation may provide insights into how to improve management strategies and potentially predict future evolutionary changes in *P. nodorum* populations.

Our aims in this study were to investigate the effects of natural selection and genetic drift on quantitative traits using eight populations of *P. nodorum* sampled from naturally infected farmer’s fields around the world. We tested the hypothesis that local environmental conditions (e.g, high or low temperatures) and agricultural practices (e.g., fungicide applications) would impose directional selection on different traits of *P. nodorum.* First, we estimated the additive genetic variation for colony growth and melanization phenotypes following exposure to a range of different temperatures (providing a measure of thermal sensitivity) and fungicide concentrations (providing a measure of fungicide sensitivity). We accomplished this by measuring colony growth rates and melanization for 164 strains of *P. nodorum* using automated image analysis (Lendenmann et al., 2014, 2015). Next, we determined the degree of population genetic structure among the eight populations using nearly 50,000 neutral SNPs extracted from whole-genome sequences for all 164 strains. Finally, we estimated Q_ST_ values for each trait and compared these values to the F_ST_ index calculated across the eight populations. The Q_ST_ - F_ST_ comparisons allowed us to infer the predominant evolutionary forces driving quantitative trait divergence among these populations and allowed us to detect local adaptation in response to high temperatures and fungicide exposure.

## 2. MATERIAL AND METHODS

### 2.1. Fungal populations and preparation of inoculum used for phenotyping

*P. nodorum* strains were sampled between 1991 and 2005 from eight wheat fields growing in eight locations, including Australia, Iran, New York (USA), Oregon (USA), Texas (USA), South Africa and Switzerland (sites A and B). Details regarding sampling sites and population genetic structure based on SSR markers were described previously (McDonald et al., 2012; Stukenbrock et al., 2006). A total of 164 genetically distinct isolates were analyzed with an average of ~21 isolates per geographical field population (Table 1).

**Table 1.** Origin, location, year of collection and sample size (*N*) of *Parastagonospora nodorum* populations included in the Q_st_-F_st_ analysis.

| Origin | Location | Year | *N* | Collector(s) |
| --- | --- | --- | --- | --- |
| *Oceania* |  |  |  |  |
| Australia | Narrogin | 2001 | 22 | B.A. McDonald & R. Loughman |
| *Africa* |  |  |  |  |
| South Africa | Southwestern Cape | 1995 | 22 | P. Crous |
| *Europe* |  |  |  |  |
| Switzerland A | Winterthur | 1999 | 22 | B.A. McDonald & V. Michel |
| Switzerland B | Bern | 1999 | 22 | B.A. McDonald & V. Michel |
| *Asia* |  |  |  |  |
| Iran | Golestan Province | 2005/2010 | 16 | R. Sommerhalder & M. Razavi |
| *North America* |  |  |  |  |
| New York | Ithaca | 1991 | 21 | G. Bergstrom |
| Oregon | Hyslop | 1993 | 17 | M. Schmidt |
| Texas | Overton | 1992 | 22 | B.A. McDonald & L. Nelson |
| Totals |  | 1992-2010 | 164 |  |

In earlier publications (McDonald et al., 2013, 2012; Pereira et al., 2017; Sommerhalder et al., 2006; Stukenbrock et al., 2006), the Switzerland 1999B population was indicated to originate from China. As a result of the genome sequence analyses reported in this paper, we believe that a transcription error led to mislabeling of the China 2001 population, which we now believe was collected in 1999 from a Swiss field of wheat located near Bern, ~150 km away from where the Swiss 1999A population was collected. The transcription error may have resulted from the fact that the isolates from China labelled CHI01 (CHI for China, 01 for 2001) were mistakenly replaced by Swiss isolates labelled CH1 (CH1, Swiss collection 1, made in 1999). We discovered this error after our genome-wide analyses revealed that the Swiss 1999 population was virtually indistinguishable from the China 2001 population based on comparison of 49374 SNP markers distributed across the genome. While this mix-up is embarrassing, it does not compromise any of the analyses or interpretations reported in this manuscript.

Based on preliminary experiments, we used 5-mm-diameter plugs of mycelium as initial inoculum for all experiments. All isolates were retrieved from −80°C long-term storage on silica gel and transferred to Petri dishes containing Potato Dextrose Agar (PDA, 4 g L^-1^ potato starch, 20 g L^-1^ dextrose, 15 g L^-1^ agar and 50 mg L^-1^ kanamycin). The PDA plates were grown for three days in the dark at a constant temperature of 24°C. 5-mm-diameter plugs of mycelium were cut from the edges of the growing colonies using a sterilized cork borer and placed onto the center of fresh PDA Petri dishes. These plates were grown in the dark at 24°C for seven days and then used as the inoculum sources for all experiments.

### 2.2. Strain phenotyping

All 164 isolates were exposed to the same seven environments, including low, optimum and high growth temperatures and four concentrations of an azole fungicide. The experiments were conducted in square Petri dishes (120 × 120 × 17 mm, Huberlab) containing PDA. Four 5-mm-diameter mycelium plugs of each strain were placed into the corners of a square plate with equidistant separation. Each treatment was replicated twice, generating eight colonies in total for each strain.

For the thermal response experiment, all isolates were grown in the dark on PDA at 18°C, 24°C and 30°C. Based on the outcomes of preliminary experiments using a subset of 16 of the strains (two from each field population), 18°C and 30°C were chosen to represent stressful temperatures while 24°C was chosen as an optimum temperature for growth. For the fungicide stress experiment, all isolates were grown at 24°C on PDA amended with propiconazole (Syngenta, Basel, Switzerland) at either 0, 0.1, 0.5 or 1 ppm. All inoculation procedures were performed on the same day for each of three separate batches of approximately 54 isolates. No significant differences could be attributed to batch effects.

Digital images for each environment were taken at 2, 4, 6 and 8 days after inoculation (DAI), a total of four time points. All camera settings and configurations, plate orientations, and lighting conditions were standardized as described previously (Lendenmann et al., 2014). After acquiring images, the plates were returned to their growth chambers and their positions in the growth chamber were re-randomized. The images were automatically analyzed using a modified version of a batch macro developed for ImageJ (Lendenmann et al., 2014). For the new macro, the conversion from pixels to square millimeters was performed using a calibration image taken from 50 cm above the Petri dish. Colony detection was performed using the color threshold option, with the hue sliding scales varying between 22 and 255 (macro lines 70 and 71) (Supplementary 1).

Quantitative measurements were acquired from image analyses for each colony. Total colony area (mm^2^) and mean grey value per colony (GV, a proxy for total melanization on the 0-255 grey scale, where 0 is completely black and 255 is completely white) were measured from digital images taken through the Petri dish top for colony size measures and the Petri dish bottom for GV measures. We obtained a total of 8 raw data points per isolate at each time point in each environment. The raw measures of total colony area and GV were used to determine the following traits: (i) Radial growth rate was obtained by fitting the mean colony radii 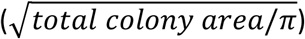 over the four time points using a general linear model, resulting in average r^2^ > 0.9 (Trinci 1971; Lendenmann et al. 2015); relative growth rates reflecting (ii) fungicide resistance and (iii) temperature sensitivity (TS) were determined for each isolate as the ratio between growth rates under different fungicide concentrations compared to the absence of fungicide, and growth rates at 18°C or 30°C compared to 24°C; (iv) melanization rate (MRate) was obtained by fitting the distribution of GVs over the four time points using a general linear model. A positive value for MRate indicates a decrease in melanization and a negative value for MRate indicates an increase in colony melanization over time. The (v) melanization response (MResp) trait was determined as the ratio between GV under a given fungicide dose and in the absence of fungicide; and as the ratio between GV at 18°C or 30°C against 24°C. When MResp > 1 it represents a decrease in melanization and MResp < 1 represents an increase in melanization after stress exposure relative to optimum conditions. Variation among isolates for GV reached its maximum at 8 DAI, so MResp was calculated based on this time point.

### 2.3. Strain genotyping

Entire genome sequences were generated for all 164 strains. Strains were grown in Potato Dextrose Broth (PDB) and total DNA was extracted from lyophilized mycelium using DNeasy Plant Mini Kits (Qiagen) according to the manufacturer’s instructions. Whole-genome sequencing was performed on an Illumina HiSeq 2500 platform, with paired-end reads of 150 bp. All the Illumina sequence data are available in the NCBI Short Read Archive (BioProject PRJNA606320).

The generated raw reads were trimmed for remaining Illumina adapters and read quality with Trimmomatic v0.36 (Bolger et al., 2014), using the following settings: illuminaclip = TruSeq3-PE.fa:2:30:10; leading = 10; trailing = 10; slidingwindow = 5:10; minlen = 50. Trimmed reads were aligned with the reference isolate SN2000 (Richards et al., 2017), which is assembled into chromosomes. The alignment was performed with the short-read aligner Bowtie 2 version 2.3.3 (Langmead and Salzberg, 2012), using the --very-sensitive-local option. Duplicated PCR reads were marked as duplicate using Picard tools version 2.17.2 (http://broadinstitute.github.io/picard).

Single nucleotide polymorphism (SNP) calling and variant filtration were performed using the Genome Analysis Toolkit (GATK) version 3.8-0 (McKenna et al., 2010). First, we used HaplotypeCaller in each isolate file individually, with the - emitRefConfidence GVCF and -ploidy 1 options. Then, joint variant calls were performed using GenotypeGVCFs with the flag-maxAltAlleles 2. Finally, SelectVariants and VariantFiltration were used for hard filtering SNPs with the following cut-offs: QUAL < 200; QD < 10.0; MQ < 20.0; –2 > BaseQRankSum > 2; – 2 > MQRankSum > 2; –2 > ReadPosRankSum > 2; FS > 0.1.

We retained only bi-allelic sites and excluded sites with missing data using vcftools 0.1.15 (Danecek et al., 2011). Using the function --indep-pairwise in plink v1.9 we pruned SNPs above a linkage disequilibrium threshold of 0.2 using a sliding window of 15 kb (Chang et al., 2015). From this unlinked SNP dataset, we selected only SNPs causing synonymous substitutions (on four-fold degenerated sites) to identify neutral SNP markers, using the software VCF2MK (https://github.com/russcd/vcf2MK). The final data set consisted of 49374 neutral and un-linked SNPs.

### 2.4. Data Analyses

We applied a general linear model to determine whether there were significant effects for populations and isolates nested within populations on the trait values (package “lm” and function anova in R) (R Core Team, 2019). Among-population comparisons of mean growth rate and mean melanization in the different environments were based on Tukey’s honest significant difference test using R.

Within-population components of variance for each trait were determined using genetic variance and heritability (Willi et al., 2011; Stefansson et al., 2014; Pereira et al., 2016). The variance components were determined using a statistical procedure implemented in R software using the lme4 package (Bates et al., 2015). The variance derived from the isolates within each population was interpreted as genetic variance (V_G_), while variance among replicates of the same clone was interpreted as environmental variance because replicates had the same genotype. Narrow-sense heritability (h^2^) was calculated as the ratio between V_G_ and total phenotypic variation within a population (Falconer and Mackay, 1996). Confidence intervals were estimated by applying a bootstrap with 999 re-samples.

Estimates of population divergence (F_ST_) were calculated using the 49374 retained SNPs with the R package hierfstat (Goudet, 2005). Overall and among-population values of F_ST_ as well as their confidence intervals were determined by bootstrapping with 999 resamplings using hierfstat (Goudet, 2005). Nei’s diversity was calculated using the popgenome R package (Pfeifer et al., 2014).

*P. nodorum* is a haploid organism, so dominance effects among alleles within loci can be ignored. If we assume a small or negligible epistatic effect within populations, genetic variance is equivalent to additive genetic variance for the determination of population divergence in quantitative traits (Q_ST_) (Whitlock, 2008). Under this scenario, the following formula can be used to calculate Q_ST_ as described by Zhan et al. (2005):

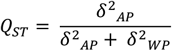

Where *δ^2^_AP_* is the additive genetic variance attributed to among-population variation and *δ^2^_WP_* is the additive genetic variance attributed to within-population variation.

Correlation analyses among the 12 traits used in the Q_ST_ - F_ST_ analysis, between overall heritability and Nei’s diversity, between MRate and fungicide resistance/growth rate and between MResp and growth rate and MRate were performed using a general linear model based on Pearson’s coefficient in the R package RcmdrMisc (Fox, 2005), and visually represented using the R package ggplot2 (Wickham, 2009).

The thermal reaction norm of *P. nodorum* was modelled based on a second-order polynomial equation using individual measures of average growth rate for every isolate across the three tested temperatures. Another measure, the composite reaction norm, was calculated based on the slope of the reaction norm between 24°C and 18°C and between 24°C and 30°C. An average value was calculated from the two slopes and populations were compared based on these isolate mean values.

## 3. RESULTS

### 3.1. Colony growth rate and melanization show quantitative distributions and high heritabilities

Phenotypic variation for traits related to growth and melanization in *P. nodorum* was assessed after exposure to different temperatures and different concentrations of an azole fungicide for eight geographical field populations (Figure 1). As expected, the environment of 24°C without fungicide, hereafter referred to as the control environment, provided the fastest growth rate for most isolates. We found near-normal distributions for growth rate and melanization in the control environment and at 18°C and 30°C, consistent with quantitative inheritance of these traits (Figure 1). Growth rates at 0.1, 0.5 and 1 ppm fungicide showed bimodal distributions (Figure 1). In general, all environments significantly affected the average trait values across all populations and across all isolates within populations (*P* ≤ 0.0001, Table 2).

**Figure 1.**
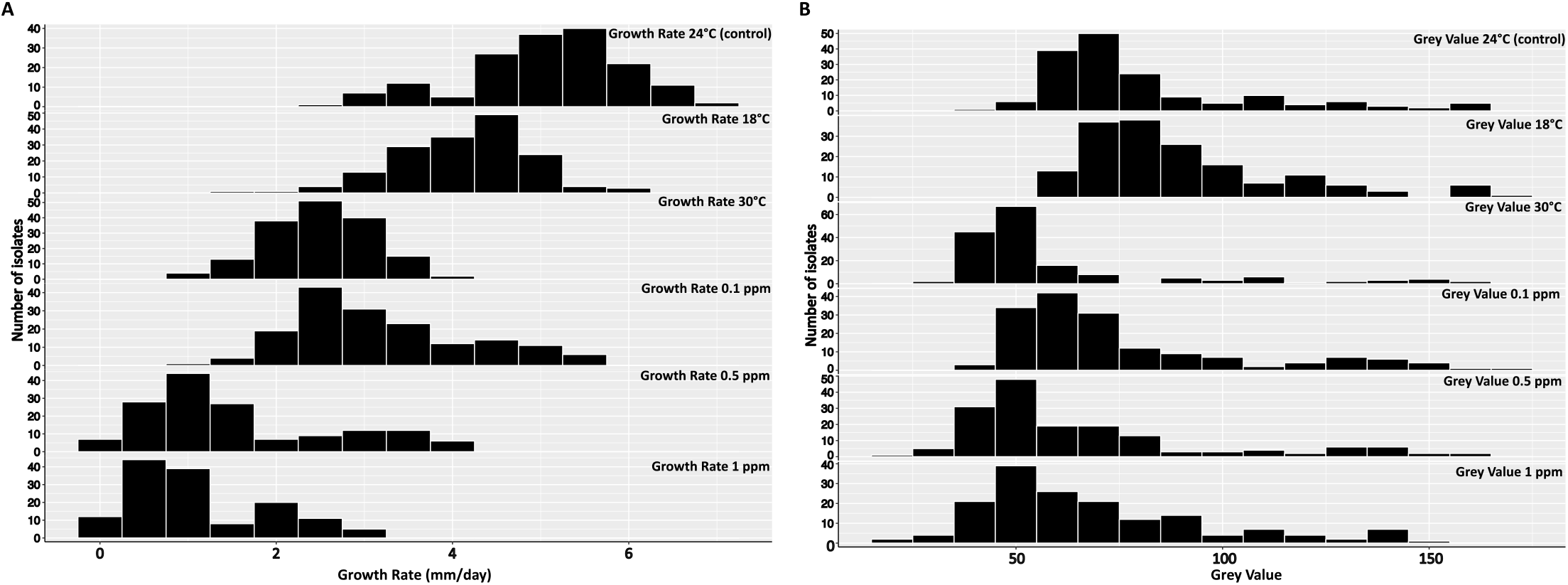
The distribution of growth rate (A) and grey value (B) for 164 isolates of *Parastagonospora nodorum* growing under different temperatures and fungicide concentrations. Each plot within a panel represents a different temperature or fungicide treatment as indicated in the aligned text. The grey value per colony is a proxy for total melanization on the 0-255 grey scale, where 0 is completely black (or highly melanized) and 255 is completely white (or not melanized).

**Table 2.** General linear model analyses testing the effect of population and isolate (nested within population) on quantitative traits of *Parastagonospora nodorum*.

| Trait | Source | df | MS | F | P |
| --- | --- | --- | --- | --- | --- |
| Growth Rate 24°C (control) | Population | 7 | 4.22 | 77.08 | <0.0001 |
|  | Isolates within population | 156 | 1.45 | 26.52 | <0.0001 |
|  | Error | 164 | 0.05 | NA | NA |
| Growth Rate 18°C | Population | 7 | 2.41 | 54.10 | <0.0001 |
|  | Isolates within population | 156 | 1.17 | 26.27 | <0.0001 |
|  | Error | 164 | 0.04 | NA | NA |
| Growth Rate 30°C | Population | 7 | 2.40 | 76.40 | <0.0001 |
|  | Isolates within population | 156 | 0.70 | 22.32 | <0.0001 |
|  | Error | 164 | 0.03 | NA | NA |
| Growth Rate 0.1 ppm | Population | 7 | 31.16 | 620.08 | <0.0001 |
|  | Isolates within population | 156 | 0.71 | 14.04 | <0.0001 |
|  | Error | 164 | 0.05 | NA | NA |
| Growth Rate 0.5 ppm | Population | 7 | 40.11 | 640.44 | <0.0001 |
|  | Isolates within population | 156 | 0.82 | 13.03 | <0.0001 |
|  | Error | 164 | 0.06 | NA | NA |
| Growth Rate 1 ppm | Population | 7 | 23.66 | 579.90 | <0.0001 |
|  | Isolates within population | 156 | 0.53 | 12.88 | <0.0001 |
|  | Error | 164 | 0.04 | NA | NA |
| Grey Value 24°C (control) | Population | 7 | 6712.08 | 52.06 | <0.0001 |
|  | Isolates within population | 156 | 1553.06 | 12.05 | <0.0001 |
|  | Error | 164 | 128.92 | NA | NA |
| Grey Value 18°C | Population | 7 | 3048.37 | 39.32 | <0.0001 |
|  | Isolates within population | 156 | 1068.70 | 13.78 | <0.0001 |
|  | Error | 164 | 77.53 | NA | NA |
| Grey Value 30°C | Population | 7 | 5365.80 | 345.83 | <0.0001 |
|  | Isolates within population | 156 | 1465.98 | 94.48 | <0.0001 |
|  | Error | 164 | 15.52 | NA | NA |
| Grey Value 0.1 ppm | Population | 7 | 5190.23 | 84.33 | <0.0001 |
|  | Isolates within population | 156 | 1693.65 | 27.52 | <0.0001 |
|  | Error | 164 | 61.55 | NA | NA |
| Grey Value 0.5 ppm | Population | 7 | 4181.42 | 9.85 | <0.0001 |
|  | Isolates within population | 156 | 1710.79 | 4.03 | <0.0001 |
|  | Error | 164 | 424.37 | NA | NA |
| Grey Value 1 ppm | Population | 7 | 4008.38 | 10.36 | <0.0001 |
|  | Isolates within population | 156 | 1402.31 | 3.62 | <0.0001 |
|  | Error | 164 | 386.93 | NA | NA |

We observed high heritability (h^2^) values for growth rate and melanization in the different environments (Table 3). h^2^ ranged from a low of 0.58 for melanization in 0.5 ppm propiconazole to a high of 0.95 for growth rate in the control environment and at 18°C and 30°C. Other traits with h^2^ higher than 0.9 were growth rates in 0.1 ppm, 0.5 ppm, and 1 ppm propiconazole and melanization in the control environment, 18°C, 30°C and in 0.1 ppm propiconazole.

**Table 3.** Heritability measures for 12 quantitative traits of *Parastagonospora nodorum*.

| Population | Growth Rate<br>24°C (control) | Growth Rate<br>18°C | Growth Rate<br>30°C | Growth Rate<br>0.1 ppm | Growth Rate<br>0.5 ppm | Growth Rate<br>1 ppm | Grey Value<br>24°C (control) | Grey Value<br>18°C | Grey Value<br>30°C | Grey Value<br>0.1 ppm | Grey Value<br>0.5 ppm | Grey Value<br>1 ppm |
| --- | --- | --- | --- | --- | --- | --- | --- | --- | --- | --- | --- | --- |
| Australia | 0.96 | 0.92 | 0.99 | 0.98 | 0.82 | 0.79 | 0.96 | 0.97 | 0.98 | 0.97 | 0.39 | 0.49 |
|  | 0.02 | 0.05 | 0.01 | 0.01 | 0.09 | 0.11 | 0.02 | 0.02 | 0.01 | 0.02 | 0.32 | 0.45 |
| South Africa | 0.99 | 0.91 | 0.96 | 0.63 | 0.89 | 0.92 | 0.98 | 0.51 | 0.59 | 0.60 | 0.44 | 0.97 |
|  | 0.01 | 0.08 | 0.03 | 0.31 | 0.07 | 0.04 | 0.01 | 0.49 | 0.39 | 0.37 | 0.34 | 0.02 |
| Switzerland A | 0.93 | 0.98 | 0.79 | 0.97 | 0.98 | 0.97 | 0.98 | 0.98 | 0.98 | 0.99 | 0.92 | 0.69 |
|  | 0.03 | 0.01 | 0.21 | 0.02 | 0.01 | 0.02 | 0.01 | 0.01 | 0.00 | 0.01 | 0.07 | 0.31 |
| Switzerland B | 0.97 | 0.95 | 0.99 | 0.94 | 0.98 | 0.96 | 0.99 | 0.99 | 0.98 | 0.96 | 0.98 | 0.99 |
|  | 0.01 | 0.02 | 0.00 | 0.03 | 0.01 | 0.02 | 0.00 | 0.00 | 0.01 | 0.02 | 0.01 | 0.01 |
| Iran | 0.99 | 0.92 | 0.99 | 0.95 | 0.93 | 0.92 | 0.98 | 0.99 | 0.97 | 0.98 | 0.50 | 0.48 |
|  | 0.00 | 0.06 | 0.00 | 0.02 | 0.05 | 0.04 | 0.00 | 0.01 | 0.00 | 0.01 | 0.33 | 0.39 |
| New York | 0.98 | 0.98 | 0.96 | 0.99 | 0.77 | 0.86 | 0.66 | 0.99 | 0.98 | 0.97 | 0.52 | 0.66 |
|  | 0.01 | 0.01 | 0.04 | 0.00 | 0.19 | 0.08 | 0.35 | 0.00 | 0.00 | 0.01 | 0.44 | 0.32 |
| Oregon | 0.77 | 0.93 | 0.98 | 0.92 | 0.98 | 0.94 | 0.98 | 0.99 | 0.98 | 0.98 | 0.47 | 0.34 |
|  | 0.18 | 0.06 | 0.01 | 0.06 | 0.01 | 0.02 | 0.00 | 0.00 | 0.00 | 0.01 | 0.39 | 0.34 |
| Texas | 0.98 | 0.98 | 0.97 | 0.97 | 0.93 | 0.96 | 0.99 | 0.99 | 0.97 | 0.89 | 0.45 | 0.45 |
|  | 0.01 | 0.01 | 0.01 | 0.01 | 0.03 | 0.02 | 0.00 | 0.00 | 0.01 | 0.06 | 0.47 | 0.47 |
| Overall Mean | 0.95 | 0.95 | 0.95 | 0.92 | 0.91 | 0.91 | 0.94 | 0.93 | 0.93 | 0.92 | 0.58 | 0.63 |
| Standard deviations | 0.07 | 0.03 | 0.07 | 0.12 | 0.08 | 0.06 | 0.11 | 0.17 | 0.14 | 0.13 | 0.23 | 0.24 |
| Coefficient of variation | 0.08 | 0.03 | 0.07 | 0.13 | 0.09 | 0.07 | 0.12 | 0.18 | 0.15 | 0.14 | 0.40 | 0.38 |

### 3.2. Extreme temperatures and fungicide exposure reduced growth rates

In the optimal control environment, the Australian population had a significantly lower growth rate (4.37 mm/day) than all other populations except South Africa (*P* ≤ 0.05, Table 4). Average growth rates slowed as propiconazole concentrations increased (*P* ≤ 0.001). At 0.1 ppm, the Chinese and Swiss populations had the fastest growth rates (4.6 and 4.3 mm/day, respectively; *P* ≤ 0.05; Table 4), while the populations from Australia, Oregon and South Africa had the slowest growth rates (2.6, 2.4 and 2.5 mm/day, respectively; *P* ≤ 0.05; Table 4). At the highest propiconazole concentrations (0.5 and 1 ppm), the Chinese population grew the fastest (3.3 and 2.4 mm/day, respectively; *P* ≤ 0.05; Table 4), followed by Switzerland (2.7 and 1.9 mm/day, respectively; *P* ≤ 0.05; Table 4), with both populations showing significantly lower fungicide sensitivity than the other populations. No differences were detected among the other populations at these two concentrations.

**Table 4.** Nei’s diversity for 12 quantitative traits of *Parastagonospora nodorum*.

| Population | Nei's Diversity | Growth Rate<br>24°C (control) | Growth Rate<br>18°C | Growth Rate<br>30°C | Growth Rate<br>0.1 ppm | Growth Rate<br>0.5 ppm | Growth Rate<br>1 ppm | Grey Value<br>24°C (control) | Grey Value<br>18°C | Grey Value<br>30°C | Grey Value<br>0.1 ppm | Grey Value<br>0.5 ppm | Grey Value<br>1 ppm |
| --- | --- | --- | --- | --- | --- | --- | --- | --- | --- | --- | --- | --- | --- |
| Australia | 0.091E* | 4.37B | 3.63B | 2.48A | 2.57CDE | 0.96C | 0.59C | 70.40C | 81.73B | 51.40C | 66.01BC | 50.88C | 56.79B |
| South Africa | 0.095DE | 4.86AB | 3.94AB | 1.93B | 2.52DE | 1.01C | 0.58C | 80.64ABC | 89.87AB | 59.43ABC | 82.19AB | 71.85ABC | 85.30A |
| Switzerland A | 0.120B | 5.26A | 4.26A | 2.52A | 4.33A | 2.66B | 1.88B | 79.74BC | 89.40AB | 57.98BC | 85.01AB | 72.11AB | 67.78AB |
| Switzerland B | 0.121B | 5.11A | 4.20A | 2.56A | 4.66A | 3.27A | 2.38A | 71.81C | 82.28B | 50.57C | 73.35ABC | 64.19ABC | 59.88B |
| Iran | 0.131A | 5.37A | 4.28A | 2.62A | 2.97BC | 1.09C | 0.65C | 94.56AB | 100.05A | 77.31A | 75.29ABC | 69.38ABC | 64.42AB |
| New York | 0.100CD | 5.25A | 4.37A | 2.49A | 2.83BCD | 0.86C | 0.51C | 93.37AB | 103.55A | 72.63AB | 84.46AB | 73.26AB | 74.82AB |
| Oregon | 0.105C | 5.11A | 4.19A | 2.57A | 2.38E | 0.65C | 0.48C | 97.43A | 97.86AB | 74.11AB | 90.80A | 77.70A | 75.04AB |
| Texas | 0.102C | 5.17A | 4.20A | 2.70A | 3.12B | 0.97C | 0.53C | 69.79C | 82.21B | 47.24C | 56.96C | 51.95BC | 60.61B |
| Overall Mean | 0.11 | 5.06 | 4.13 | 2.48 | 3.17 | 1.44 | 0.95 | 82.22 | 90.87 | 61.33 | 76.76 | 66.42 | 68.08 |
| Standard deviations | 0.01 | 0.32 | 0.24 | 0.24 | 0.86 | 0.97 | 0.74 | 11.46 | 8.70 | 11.80 | 11.18 | 10.00 | 9.67 |

\*Values followed by different letters in the same column are significantly different at $P \leq 0.05$ .

Temperatures of 18°C and 30°C significantly reduced the average growth rates compared to the control temperature (*P* ≤ 0.001). At 18°C, the fastest growth rate was in the New York population, and the slowest was in Australia (4.4 and 3.6 mm/day, respectively, *P* ≤ 0.05, Table 4). At 30°C, there were no significant differences among populations except for South Africa, which had the slowest growth rate (1.9 mm/day, *P* ≤ 0.05, Table 4). Based on the TS calculations, the populations were overall more sensitive to higher temperatures than to lower temperatures (Supplementary 2). For TS at the higher temperature, the populations from Australia and Texas were the least sensitive (TS values were closest to 1), whereas the population from South Africa was the most sensitive. For TS at the lower temperature, the populations from New York and Australia were the least affected. Altogether, 15 isolates had TS > 1 at the lower temperature and no isolate had TS > 1 at the higher temperature.

The thermal reaction norm of *P. nodorum,* which reflects the patterns of growth rate across the tested temperatures, showed a good fit to a second-order polynomial model (Supplementary 3). On average, this model accounted for 66% of the total variation for growth rates in *P. nodorum (P* ≤ *0.001).* We did not find significant differences among populations using this composite reaction norm.

### 3.3. Melanization varied according to environmental conditions

We observed significant variation in melanization at the population level across varying temperatures and fungicide concentrations. In the control environment, the strains in the population from Oregon showed significantly lower melanization (i.e. higher GVs) than the populations from Australia, Switzerland (1999A and 1999B) and Texas (*P* ≤ 0.05; Table 4). On average, more stressful temperatures and fungicide stress significantly affected melanization (*P* ≤ 0.001).

Melanization was significantly higher at 0.5 ppm than at 0 and 0.1 ppm of propiconazole across all populations (*P* ≤ 0.001), with mean GVs of 66, 82 and 77, respectively (Table 4). Average melanization across populations at 1 ppm (GV = 68) was not significantly different from melanization at 0.5 ppm (Table 4). Texas was the most melanized population at 0.1 ppm, while Australia was the most melanized population at 0.5 and 1 ppm propiconazole (*P* ≤ 0.05, Table 4).

Across all populations we observed higher melanization at 30°C and lower melanization at 18°C (mean GVs of 61 and 90 respectively, *P* ≤ 0.001, Table 4). At the highest temperature, Texas was the most melanized population and at the lowest temperature Australia showed the highest amount of melanization (mean GVs of 47 and 81 respectively).

### 3.4. Low population structure based on neutral genome-wide SNPs

We inferred the population genetic structure among populations using markers distributed across the entire genome. We calculated the genome-wide F_ST_ using 49374 unlinked SNPs (Supplementary 4). The overall F_ST_ across populations was 0.12 (*P* ≤ 0.0001). The pairwise F_ST_ ranged from 0.24 between Australia and Iran to no differentiation between Switzerland A and B (Table 5). Iran was the population with the highest overall Nei’s diversity (Table 4, *P* ≤ 0.05, 0.13), consistent with previous studies placing the *P. nodorum* center of origin in the fertile crescent (McDonald et al., 2012).

**Table 5.** Estimated pairwise F_ST_ according to Nei (1987) based on 49429 neutral SNPs from 164 isolates of *Parastagonospora nodorum.* Significance thresholds based on 20000 *bootstraps: *P* < 0.01, **Nonsignificant *P* ≤ 0.05.

|  | Australia | South Africa | Switzerland A | Switzerland B | Iran | New York | Oregon | Texas |
| --- | --- | --- | --- | --- | --- | --- | --- | --- |
| Australia |  | 0.16 | 0.13 | 0.12 | 0.24 | 0.17 | 0.15 | 0.18 |
| South Africa | * |  | 0.15 | 0.15 | 0.22 | 0.20 | 0.19 | 0.20 |
| Switzerland A | * | * |  | 0.00 | 0.15 | 0.08 | 0.06 | 0.08 |
| Switzerland B | * | * | **NS |  | 0.15 | 0.08 | 0.06 | 0.08 |
| Iran | * | * | * | * |  | 0.20 | 0.20 | 0.20 |
| New York | * | * | * | * | * |  | 0.08 | 0.04 |
| Oregon | * | * | * | * | * | * |  | 0.08 |
| Texas | * | * | * | * | * | * | * |  |

### 3.5. Natural selection was the predominant evolutionary force acting on P. nodorum populations

To disentangle the effects of natural selection and genetic drift on quantitative traits in populations of *P. nodorum,* we compared the Q_ST_ index for each trait to the F_ST_ index (Figure 2). The Q_ST_ values for melanization were consistently lower than F_ST_ across all environments (Figure 2, *P* ≤ 0.0001), suggesting that melanization is under stabilizing selection. Growth rates in all environments, except for the control environment and 18°C, had significantly higher Q_ST_ than F_ST_ (Figure 2, *P* ≤ 0.0001), suggesting that selection operates to favor local adaptation for these traits.

**Figure 2.**
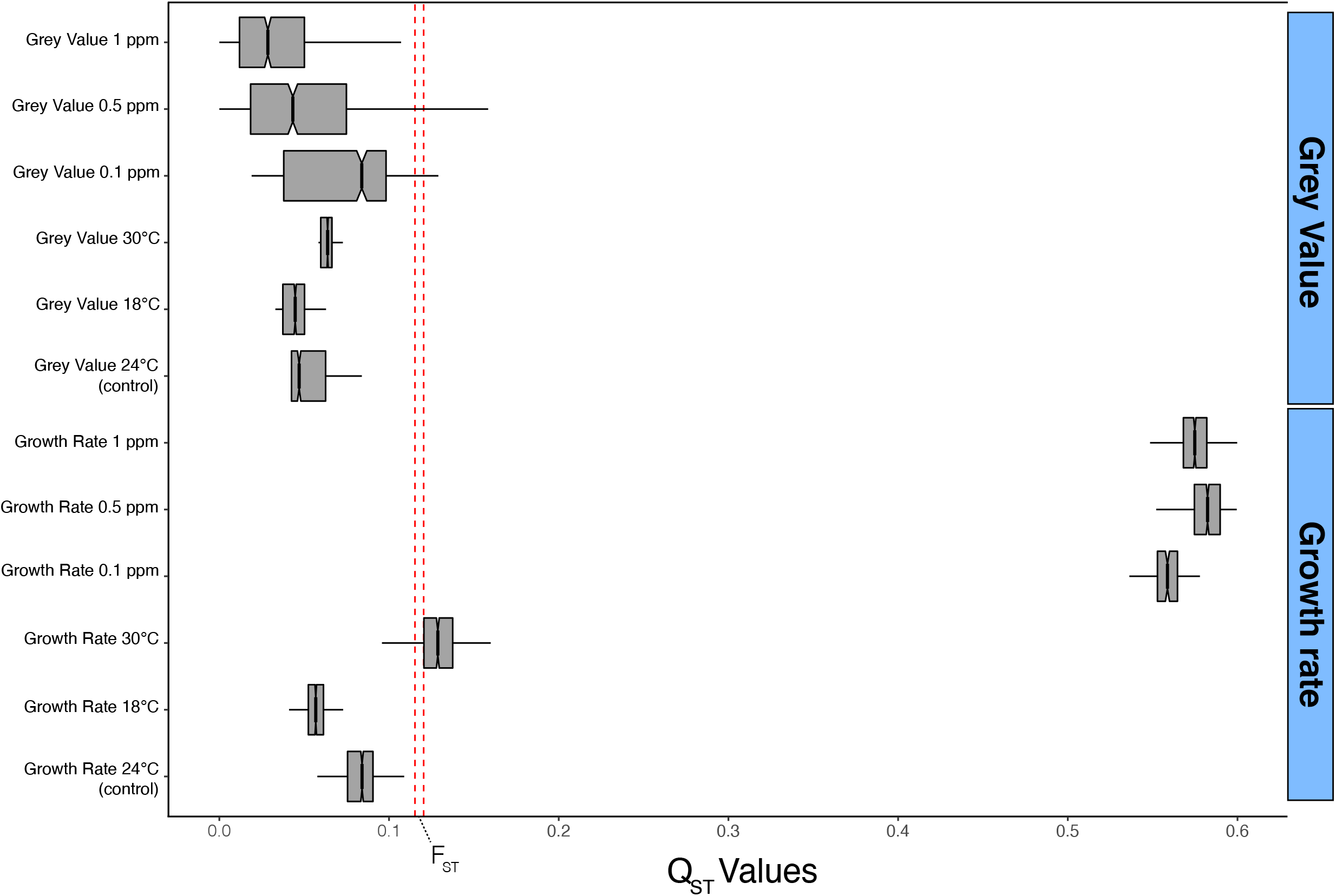
Comparisons of Q_ST_ - F_ST_ across 12 quantitative traits of *Parastagonospora nodorum.* Boxplots display the confidence intervals determined by bootstrapping with 999 resamplings of Q_ST_ values. The red dashed lines show the overall F_ST_ distribution. Traits with a Q_ST_ distribution below that range are interpreted as being under stabilizing selection, while traits with Q_ST_ distribution above that range are interpreted as being under diversifying selection. The grey value per colony is a proxy for total melanization on the 0-255 grey scale, where 0 is completely black (or highly melanized) and 255 is completely white (or not melanized).

### 3.6. Trait correlations

We next sought to investigate the relationship between pairs of traits. The correlation analysis revealed significant positive correlations among growth rates and among GVs (Figure 3, *P* ≤ 0.0001), but no significant correlations were found between growth rates and GVs for any treatment. Overall growth rates amongst the three concentrations of fungicide were all significantly correlated, indicating that less sensitive isolates maintained higher growth rates across all concentrations of propiconazole. Growth rate in the control environment was correlated with growth rate at 18°C but not with growth rate at 30°C. For melanization, we found positive correlations for GV at 0.1 ppm, 0.5 ppm, 1 ppm, 30°C, 18°C and the control environment. For instance, GV at 30°C was moderately correlated with GVs at 0.1 and 0.5 ppm propiconazole *(R* = 0.40 and 0.35 respectively, *P* ≤ 0.0001) suggesting that melanin accumulates similarly under these conditions.

**Figure 3.**
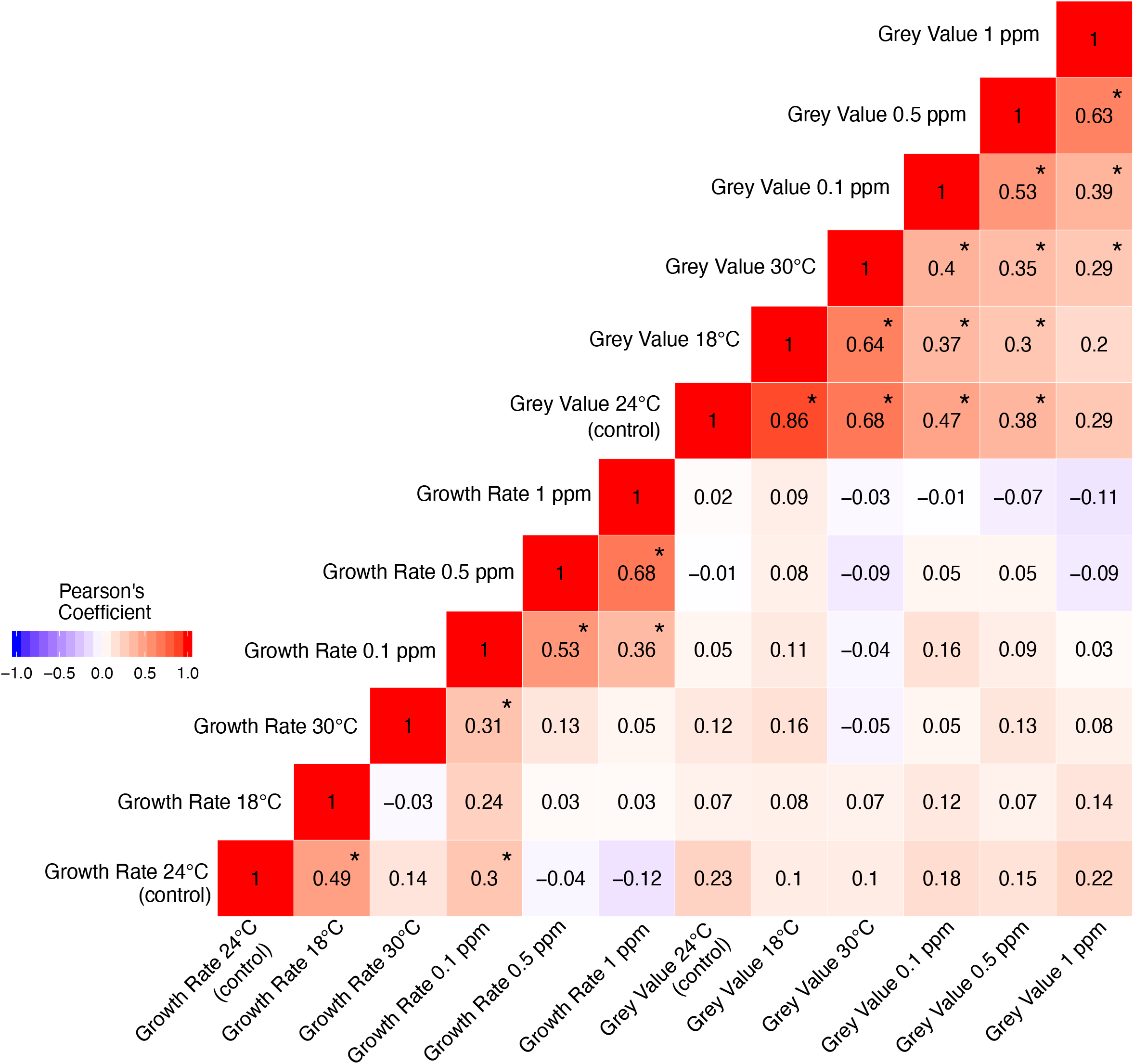
Pairwise correlations among 12 quantitative traits of *Parastagonospora nodorum.* To make data comparable across populations, trait values of each trait-population combination were standardized to a mean of 0 and a standard deviation of 1, and then correlation analyses were performed based on isolate means. Significance levels were determined after Bonferroni correction for multiple comparisons. * Significant at p < 0.0001). The grey value per colony is a proxy for total melanization on the 0-255 grey scale, where 0 is completely black (or highly melanized) and 255 is completely white (or not melanized).

Fungicide resistance and growth rates at different temperatures were significantly correlated with melanization rates (Figure 4). We found significant correlations between fungicide resistance and MRate at 0.5 and 1 ppm (*p* ≤ 0.001, Figure 4A). At the higher concentrations, isolates with more negative MRate displayed an overall increase in levels of fungicide resistance. At 30°C, growth rate and MRate were significantly correlated (Figure 4B), suggesting that isolates with higher MRate were growing faster. Analogous patterns were observed for fungicide resistance and MResp (Supplementary 5A).

**Figure 4.**
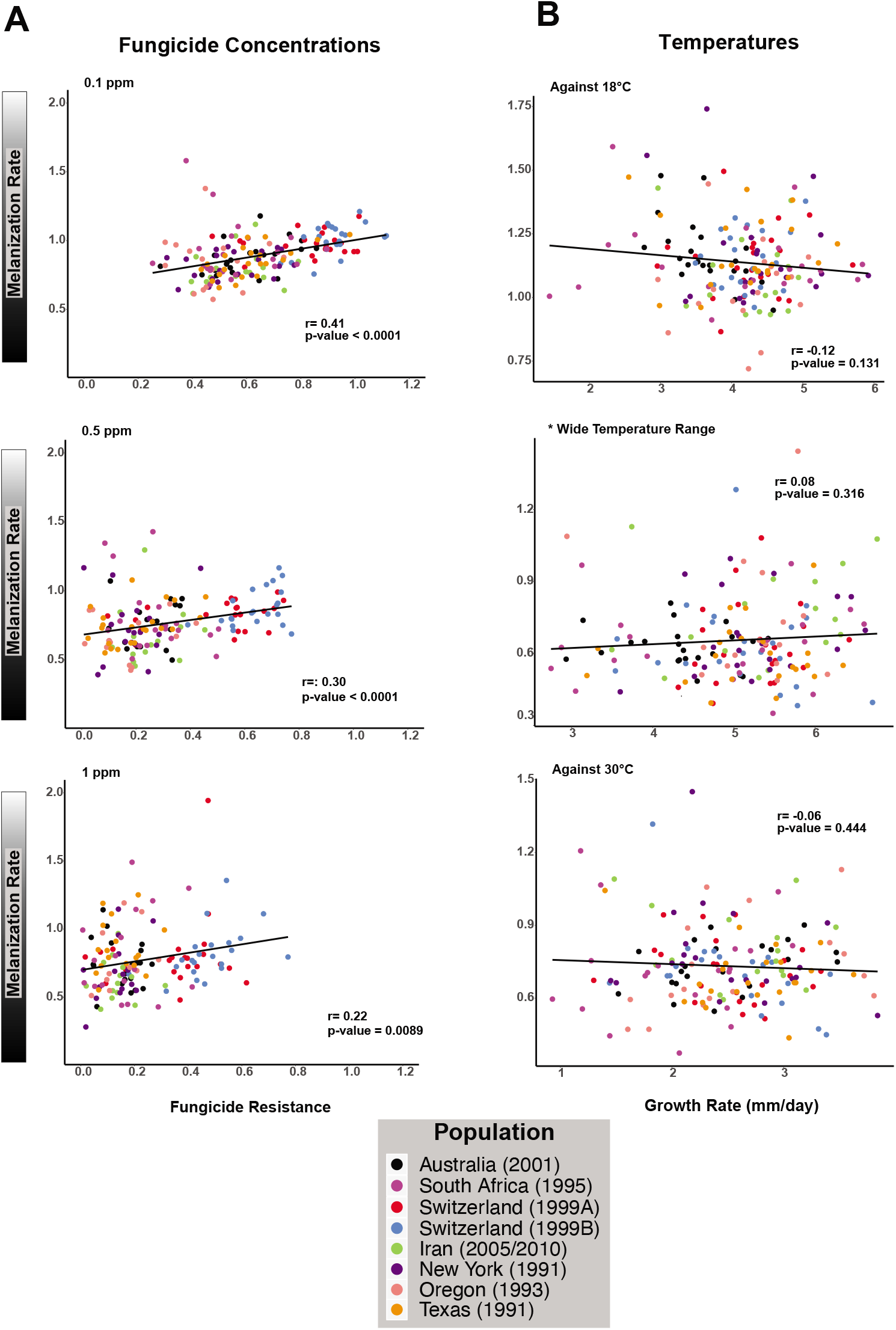
Correlation analysis between melanization rate (calculated from the slope of a line fitted to GV over time) and (A) fungicide resistance (= growth rate in presence of fungicide / growth rate in absence of fungicide) and (B) growth rate of isolates under different temperatures. The dots represent individual isolates and colours the corresponding population of origin.

No significant correlations were found between TS and mean annual temperature or annual temperature variation (Supplementary 2).

## 4. DISCUSSION

We inferred the patterns of selection operating on different quantitative traits using 164 isolates representing global field populations of the wheat pathogen *P. nodorum.* Our study supports the hypothesis that natural selection is affecting growth rates and melanization at different temperatures and fungicide concentrations, likely reflecting the process of local adaptation.

Temperature has an especially large impact on the physiology of ectothermic organisms like fungi because their internal temperature directly reflects the external thermal environment (Angilletta et al., 2006; Knies and Kingsolver, 2010). In general, phenotypic plasticity and genetic differentiation are considered the main mechanisms underlying the evolution of thermal adaptation (Chevin et al., 2010; Cooper et al., 2012; Tonsor et al., 2013; Yampolsky et al., 2013). On average, we found that plasticity made only a small contribution (<5%) to overall phenotype variation. The field populations from Australia and South Africa had both the slowest overall growth rates and the lowest levels of genetic diversity. We hypothesize that the slow growth rates in these populations reflects a lower evolutionary potential due to a founder effect (Carson 1961; Templeton et al. 2001), as also seen for Australian populations of *Z. tritici* and *R. commune* (Zhan and McDonald, 2011; Stefansson et al., 2014). The Australian and South African populations of *P. nodorum* likely originated on infected seeds when Europeans introduced wheat into these regions during the last 500 years, providing a restricted gene pool (Stukenbrock et al., 2006). Subsequent gene flow that could increase local genetic diversity during the modern era may have been prevented by global trading patterns (Australia is a major wheat exporter and South Africa imports relatively little wheat) coupled with effective quarantine measures that limited the introduction of additional infected wheat seeds or grains (Oliver et al., 2012).

The Q_ST_ - F_ST_ analyses indicated that growth at 18°C and 24°C were under stabilizing selection (Q_ST_ < F_ST_), while diversifying selection (Q_ST_ > F_ST_) was the predominant evolutionary force affecting growth at 30°C. These three temperatures represent the range of temperatures that are likely to be encountered by many *P. nodorum* populations during the wheat cropping season. It was postulated that field environments experiencing a wide fluctuation in temperatures would favor isolates that can grow more quickly and consequently colonize the host faster when conditions become conducive for disease development (Stefansson et al., 2013; Yang et al., 2016). However, because the pathogen depends on the host for reproduction, the pathogen’s overall fitness may be negatively affected if the host life span is reduced due to excessive host damage caused by pathogen growth that is too rapid (Boots et al., 2004). This trade-off between pathogen virulence and pathogen reproduction may stabilize rates of host colonization in environments where optimum temperatures for infection and colonization are more constant and frequent, offsetting the selective pressure that favors faster growers (Alizon et al., 2009). The temperatures of 18°C and 24°C used in our experiment appear closest to the optimum temperatures for *P. nodorum* growth and development, consistent with the finding of stabilizing selection, which favors isolates with growth rates closer to the population mean. The significantly higher population differentiation at 30°C is consistent with a process of diversifying selection for local adaptation. Over time, this selective process would be expected to evolve *P. nodorum* populations that are locally adapted to higher temperatures (Hayden et al., 2014; Yang et al., 2016). Given current patterns of global trade, strains of *P. nodorum* that are adapted to higher temperatures in wheat-exporting countries like Australia could be unintentionally introduced into regions such as Switzerland where local populations are maladapted to high temperatures. The long-distance movement of new strains of the wheat yellow rust fungus showing novel temperature adaptations caused extensive damage (Hovmoller et al., 2008). Thus *P. nodorum* joins the ranks of plant pathogens that are likely to pose an increasing risk to global food security in a warming world (Hovmoller et al., 2008; Milus et al., 2009; Fisher et al., 2012; Stefansson et al., 2013).

Directional selection (QST > FST) was associated with growth rates at all tested fungicide concentrations, indicating that natural selection is the main contributor to population differentiation for fungicide sensitivity. We extracted and analyzed the *CYP51* gene from the genome sequences of all 164 isolates used in this Q_ST_ - F_ST_ analysis, and confirmed the occurrence of previously reported *CYP51* mutations in isolates exhibiting the highest fungicide resistance (Pereira et al., 2017). The observed mutations, found only in the populations from Switzerland, are likely responsible for most of the differences in growth rate at different fungicide concentrations among populations.

Evidence that directional selection shapes local adaptation for fungicide resistance was also found in the wheat pathogen *Zymoseptoria tritici* (Zhan et al., 2005), while in *Rhynchosporium commune* and *Phytophthora infestans* it was found to be under stabilizing selection (Stefansson et al., 2014; Qin et al., 2016). In *Z. tritici* the emergence of *CYP51* mutations in the pathogen populations was proposed to occur locally and then spread via gene flow across Europe or, as shown more recently, across Tasmania (Brunner et al., 2008; McDonald et al., 2018). *P. nodorum* and *Z. tritici* frequently coinfect wheat plants in the field (Gilbert and Woods, 2001; Blixt et al., 2010; Oliver et al., 2012). Though the majority of fungicides applied to wheat in Europe target *Z. tritici* (Fones and Gurr, 2015), we hypothesize that these treatments indirectly selected for fungicide resistance in populations of *P. nodorum* (Knorr et al., 2019). The findings of CYP51 mutations associated with azole resistance, high heritability, and evidence for diversifying selection favoring local adaptation suggest a significant risk for emergence and spread of azole resistance over larger geographical scales for *P. nodorum.*

Melanin is a secondary metabolite found broadly across eukaryotes that often displays complex phenotypic variation across an organism’s life cycle (Butler and Day, 1998; Chumley and Valent, 1990; Dadachova and Casadevall, 2008; Singaravelan et al., 2008; Sturm and Duffy, 2012). The ecological roles associated with fungal melanin vary widely among species (Butler and Day, 1998), but it is most often reported to be related to virulence, competition with other microbes, protection against UV light and toxic compounds, and tolerance of extreme temperatures (Nosanchuk and Casadevall, 2003; Dadachova et al., 2007; Hagiwara et al., 2017). Although melanin is thought to provide protection against cold and heat stress (Rehnstrom and Free, 1996; Paolo et al., 2006), there are exceptions (Wheeler and Bell, 1988), and information regarding its impact on fungal thermal tolerance remains scarce (Cordero and Casadevall, 2017). Melanin production is energetically costly (Calvo et al., 2002), and may impose a fitness penalty if it reduces growth (Choi and Goodwin, 2011; Krishnan et al., 2018).

Quantitative measures of melanization in fungal colonies were previously used to infer possible contributions of melanin to variation in temperature and fungicide sensitivity in the fungi *Z. tritici* and *R. commune* (Lendenmann et al., 2014; Stefansson et al., 2014; Lendenmann et al., 2015; Zhu et al., 2018). In *P. nodorum* we found that stabilizing selection was the predominant evolutionary force shaping differences in melanization across all tested conditions. The low differentiation among populations for melanization suggests that selection operates against extreme phenotypes in *P. nodorum* (Sanjak et al., 2018). For the barley scald pathogen *R. commune,* melanization was also found to be under stabilizing selection (Stefansson et al., 2014). In *R. commune,* higher melanization was positively correlated with higher growth rates at 18°C and 22°C, as well as increased fungicide resistance (Zhu et al., 2018). Melanization was also correlated with faster growth rates at 15°C and reduced fungicide sensitivity in *Z. tritici* (Lendenmann et al., 2014, 2015).

In *P. nodorum* the GV for most isolates was lowest at 30°C, suggesting that melanin production increases under heat stress. We observed a negative correlation between MRate and growth rate at 30°C (*P* = 0.02), with faster-growing colonies accumulating melanin at a slower rate, but there was no correlation between growth rate and MRate at 18 or 24°C. Altogether, these findings suggest a possible trade-off between melanization and growth rate under heat stress. The calculated values for both MRate and MResp suggest that the strains that were slowest to melanize had the greatest resistance to propiconazole, indicating that melanin production did not reduce azole sensitivity. In a knockout study in *Z. tritici,* albino mutants lacking the melanin-related transcription factor *Zmr1* were exposed to two different fungicides, and had their growth compared to the wild type strain producing melanin (Krishnan et al., 2018). The fungicide bixafen (a succinate dehydrogenase inhibitor, SDHI) significantly reduced the growth of the albino mutants compared to the isogenic wild type strain, but there was no difference in the growth of the strains exposed to propiconazole (Krishnan et al., 2018). The protective effect of melanin against fungicides is attributed to its binding capacity, which reduces the fungicide availability (Bridelli et al., 2006; Paolo et al., 2006; Eisenman and Casadevall, 2012). Since melanin has a low binding affinity with azoles, the inefficient binding process may limit its overall contribution to azole resistance (Nosanchuk and Casadevall, 2006), consistent with our findings in *P. nodorum.*

Our findings illustrate how local environmental conditions can couple with different agricultural practices to shape the evolutionary trajectories of geographically distinct populations of a plant pathogen. The high heritability for traits related to fungicide and thermal sensitivity indicate the importance of genetic diversity in affecting the adaptative potential of *P. nodorum.* We found that directional selection favors genotypes with faster growth rates under fungicide and high-temperature stress. This suggests a significant risk for *P. nodorum* to develop fungicide resistance. Moreover, under the expected scenario of global warming, it is likely that SNB will easily adapt to susceptible wheat crops growing in warmer areas.

## Acknowledgements

This study was financed in part by the “Coordenação de Aperfeiçoamento de Pessoal de Nível Superior - Brasil (CAPES)” - Finance Code 001. Marcello Zala provided technical assistance. The Genetic Diversity Center (GDC) – ETH Zurich and the Functional Genomics Center in Zurich provided sequencing facilities.

## References

Alizon, S., Hurford, A., Mideo, N., Baalen, M.V., 2009. Virulence evolution and the trade-off hypothesis: history, current state of affairs and the future. Journal of Evolutionary Biology 22, 245–259. https://doi.org/10.1111/j.1420-9101.2008.01658.x

Angilletta, M., Oufiero, C.E., Leaché, A.D., 2006. Direct and Indirect Effects of Environmental Temperature on the Evolution of Reproductive Strategies: An Information-Theoretic Approach. The American Naturalist 168, E123–E135. https://doi.org/10.1086/507880

Bates, D., Mächler, M., Bolker, B., Walker, S., 2015. Fitting Linear Mixed-Effects Models Using lme4. J. Stat. Soft. 67. https://doi.org/10.18637/jss.v067.i01

Blixt, E., Olson, Å., Lindahl, B., Djurle, A., Yuen, J., 2010. Spatiotemporal variation in the fungal community associated with wheat leaves showing symptoms similar to stagonospora nodorum blotch. European Journal of Plant Pathology 126, 373–386. https://doi.org/10.1007/s10658-009-9542-z

Bolger, A., Lohse, M., Usadel, B., 2014. Trimmomatic: a flexible trimmer for Illumina sequence data. Bioinformatics 30. https://doi.org/10.1093/bioinformatics/btu170

Boots, M., Hudson, P.J., Sasaki, A., 2004. Large Shifts in Pathogen Virulence Relate to Host Population Structure. Science 303, 842–844. https://doi.org/10.1126/science.1088542

Bridelli, M.G., Ciati, A., Crippa, P.R., 2006. Binding of chemicals to melanins re-examined: Adsorption of some drugs to the surface of melanin particles. Biophysical Chemistry 119, 137–145. https://doi.org/10.1016/j.bpc.2005.06.004

Brunner, P.C., Stefanato, F.L., McDonald, B.A., 2008. Evolution of the *CYP51* gene in *Mycosphaerella graminicola:* evidence for intragenic recombination and selective replacement. Molecular Plant Pathology 9, 305–316. https://doi.org/10.1111/j.1364-3703.2007.00464.x

Butler, M., Day, A., 1998. Fungal melanins: a review. Canadian Journal of Microbiology 44, 1115–1136. https://doi.org/10.1139/w98-119

Calvo, A.M., Wilson, R.A., Bok, J., Keller, N.P., 2002. Relationship between Secondary Metabolism and Fungal Development. Microbiology and Molecular Biology Reviews 66, 447–459. https://doi.org/10.1128/mmbr.66.3.447-459.2002

Chang, C.C., Chow, C.C., Tellier, L., Vattikuti, S., Purcell, S.M., Lee, J.J., 2015. Second-generation PLINK: rising to the challenge of larger and richer datasets. GigaScience 4, 1–16. https://doi.org/10.1186/s13742-015-0047-8

Chevin, L.-M., Lande, R., Mace, G.M., 2010. Adaptation, Plasticity, and Extinction in a Changing Environment: Towards a Predictive Theory. PLoS Biology 8, e1000357. https://doi.org/10.1371/journal.pbio.1000357

Choi, Y.-E., Goodwin, S.B., 2011. Gene encoding a c-type cyclin in*Mycosphaerella graminicola* is involved in aerial mycelium formation, filamentous growth, hyphal swelling, melanin biosynthesis, stress response, and pathogenicity. Molecular plantmicrobe interactions: MPMI 24, 469–77. https://doi.org/10.1094/mpmi-04-10-0090

Chumley, F.G., Valent, B., 1990. Genetic Analysis of Melanin-Deficient, Nonpathogenic Mutants of *Magnaporthe grisea*. Molecular Plant-Microbe Interactions 3, 135. https://doi.org/10.1094/mpmi-3-135

Cooper, B.S., Tharp, J.M., Jernberg, I.I., Angilletta, M.J., 2012. Developmental plasticity of thermal tolerances in temperate and subtropical populations of Drosophila melanogaster. Journal of Thermal Biology 37, 211–216. https://doi.org/10.1016/jjtherbio.2012.01.001

Cordero, R.J.B., Casadevall, A., 2017. Functions of fungal melanin beyond virulence. Fungal Biology Reviews 31, 99–112. https://doi.org/10.1016/j.fbr.2016.12.003

Corredor-Moreno, P., Saunders, D.G.O., 2019. Expecting the unexpected: factors influencing the emergence of fungal and oomycete plant pathogens. New Phytologist. https://doi.org/10.1111/nph.16007

Dadachova, E., Bryan, R.A., Huang, X., Moadel, T., Schweitzer, A.D., Aisen, P., Nosanchuk, J.D., Casadevall, A., 2007. Ionizing Radiation Changes the Electronic Properties of Melanin and Enhances the Growth of Melanized Fungi. PLoS ONE 2, e457. https://doi.org/10.1371/journal.pone.0000457

Dadachova, E., Casadevall, A., 2008. Ionizing radiation: how fungi cope, adapt, and exploit with the help of melanin. Current Opinion in Microbiology 11, 525–531. https://doi.org/10.1016/j.mib.2008.09.013

Danecek, P., Auton, A., Abecasis, G., Albers, C.A., Banks, E., DePristo, M.A., Handsaker, R.E., Lunter, G., Marth, G.T., Sherry, S.T., McVean, G., Durbin, R., Group, 1000, 2011. The variant call format and VCFtools. Bioinformatics 27, 2156–2158. https://doi.org/10.1093/bioinformatics/btr330

Eisenman, H.C., Casadevall, A., 2012. Synthesis and assembly of fungal melanin. Applied Microbiology and Biotechnology 93, 931–940. https://doi.org/10.1007/s00253-011-3777-2

Elderd, B.D., Reilly, J.R., 2014. Warmer temperatures increase disease transmission and outbreak intensity in a host-pathogen system. Journal of Animal Ecology 83, 838–849. https://doi.org/10.1111/1365-2656.12180

Falconer, D.S., Mackay, T.F., 1996. Introduction to quantitative genetics. Longman.

Ficke, A., Cowger, C., Bergstrom, G.C., Brodal, G., 2017. Understanding yield loss and pathogen biology to improve disease management: Septoria nodorum blotch - a case study in wheat. Plant Disease. https://doi.org/10.1094/pdis-09-17-1375-fe

Fisher, M.C., Henk, Daniel.A., Briggs, C.J., Brownstein, J.S., Madoff, L.C., McCraw, S.L., Gurr, S.J., 2012. Emerging fungal threats to animal, plant and ecosystem health. Nature 484, 186. https://doi.org/10.1038/nature10947

Fones, H., Gurr, S., 2015. The impact of Septoria tritici Blotch disease on wheat: An EU perspective. Fungal Genetics and Biology 79, 3–7. https://doi.org/10.1016/j.fgb.2015.04.004

Fox, J., 2005. The *R* Commander: A Basic-Statistics Graphical User Interface to *R*. J. Stat. Soft. 14. https://doi.org/10.18637/jss.v014.i09

Gilbert, J., Woods, S.M., 2001. Leaf spot diseases of spring wheat in southern Manitoba farm fields under conventional and conservation tillage. Canadian Journal of Plant Science 81, 551–559. https://doi.org/10.4141/p00-088

Goudet, J., 2005. hierfstat, a package for r to compute and test hierarchical F-statistics. Mol Ecol Notes 5, 184–186. https://doi.org/10.1111/j.1471-8286.2004.00828.x

Hagiwara, D., Sakai, K., Suzuki, S., Umemura, M., Nogawa, T., Kato, N., Osada, H., Watanabe, A., Kawamoto, S., Gonoi, T., Kamei, K., 2017. Temperature during conidiation affects stress tolerance, pigmentation, and trypacidin accumulation in the conidia of the airborne pathogen *Aspergillus fumigatus*. PLOS ONE 12, e0177050. https://doi.org/10.1371/journal.pone.0177050

Hayden, E.J., Bratulic, S., Koenig, I., Ferrada, E., Wagner, A., 2014. The Effects of Stabilizing and Directional Selection on Phenotypic and Genotypic Variation in a Population of RNA Enzymes. Journal of Molecular Evolution 78, 101–108. https://doi.org/10.1007/s00239-013-9604-x

Hovmoller, M., Yahyaoui, A., Milus, E., Justesen, A., 2008. Rapid global spread of two aggressive strains of a wheat rust fungus. Molecular Ecology 17, 3818–3826. https://doi.org/10.1111/j.1365-294x.2008.03886.x

Knies, J.L., Kingsolver, J.G., 2010. Erroneous Arrhenius: Modified Arrhenius Model Best Explains the Temperature Dependence of Ectotherm Fitness. The American Naturalist 176, 227–233. https://doi.org/10.1086/653662

Knorr, K., Jørgensen, L.N., Nicolaisen, M., 2019. Fungicides have complex effects on the wheat phyllosphere mycobiome. PLOS ONE 14, e0213176. https://doi.org/10.1371/journal.pone.0213176

Krishnan, P., Meile, L., Plissonneau, C., Ma, X., Hartmann, F., Croll, D., McDonald, B., Sánchez-Vallet, A., 2018. Transposable element insertions shape gene regulation and melanin production in a fungal pathogen of wheat. BMC Biology. https://doi.org/10.1186/s12915-018-0543-2

Laine, A., 2008. Temperature-mediated patterns of local adaptation in a natural plantpathogen metapopulation. Ecology Letters 11, 327–337. https://doi.org/10.1111/j.1461-0248.2007.01146.x

Langmead, B., Salzberg, S., 2012. Fast gapped-read alignment with Bowtie 2. Nature Methods 9. https://doi.org/10.1038/nmeth.1923

Leinonen, T., McCairns, R.J.S., O’Hara, R.B., Merilä, J., 2013. QST-FST comparisons: evolutionary and ecological insights from genomic heterogeneity. Nature Reviews Genetics 14, 179. https://doi.org/10.1038/nrg3395

Leinonen, T., O’Hara, R., Cano, J., Merilä, J., 2008. Comparative studies of quantitative trait and neutral marker divergence: a meta-analysis. Journal of Evolutionary Biology 21, 1–17. https://doi.org/10.1111/j.1420-9101.2007.01445.x

Lendenmann, M.H., Croll, D., McDonald, B.A., 2015. QTL mapping of fungicide sensitivity reveals novel genes and pleiotropy with melanization in the pathogen *Zymoseptoria tritici*. Fungal Genetics and Biology 80, 53–67. https://doi.org/10.1016/j.fgb.2015.05.001

Lendenmann, M.H., Croll, D., Stewart, E.L., McDonald, B.A., 2014. Quantitative Trait Locus Mapping of Melanization in the Plant Pathogenic Fungus *Zymoseptoria tritici*. G3: Genes|Genomes|Genetics 4. https://doi.org/10.1534/g3.114.015289

McDonald, B.A., Stukenbrock, E.H., 2016. Rapid emergence of pathogens in agroecosystems: global threats to agricultural sustainability and food security. Philosophical Transactions of the Royal Society B: Biological Sciences 371, 20160026. https://doi.org/10.1098/rstb.2016.0026

McDonald, M.C., Oliver, R.P., Friesen, T.L., Brunner, P.C., McDonald, B.A., 2013. Global diversity and distribution of three necrotrophic effectors in *Phaeosphaeria nodorum* and related species. New Phytologist 199, 241–251. https://doi.org/10.1111/nph.12257

McDonald, M.C., Razavi, M., Friesen, T.L., Brunner, P.C., McDonald, B.A., 2012. Phylogenetic and population genetic analyses of *Phaeosphaeria nodorum* and its close relatives indicate cryptic species and an origin in the Fertile Crescent. Fungal Genetics and Biology 49. https://doi.org/10.1016/j.fgb.2012.08.001

McDonald, M.C., Renkin, M., Spackman, M., Orchard, B., Croll, D., Solomon, P.S., Milgate, A., 2018. Rapid parallel evolution of azole fungicide resistance in Australian populations of the wheat pathogen *Zymoseptoria tritici*. Applied and Environmental Microbiology. https://doi.org/10.1128/AEM.01908-18

McKenna, A., Hanna, M., Banks, E., Sivachenko, A., Cibulskis, K., Kernytsky, A., Garimella, K., Altshuler, D., Gabriel, S., Daly, M., DePristo, M., 2010. The Genome Analysis Toolkit: A MapReduce framework for analyzing next-generation DNA sequencing data. Genome Research 20. https://doi.org/10.1101/gr.107524.110

Merilä, J., Crnokrak, P., 2001. Comparison of genetic differentiation at marker loci and quantitative traits. Journal of Evolutionary Biology 14, 892–903. https://doi.org/10.1046/j.1420-9101.2001.00348.x

Milus, E.A., Kristensen, K., Hovmøller, M.S., 2009. Evidence for Increased Aggressiveness in a Recent Widespread Strain of *Puccinia striiformis* f. sp. *tritici* Causing Stripe Rust of Wheat. Phytopathology 99, 89–94. https://doi.org/10.1094/phyto-99-1-0089

Murray, G., Brennan, J., 2009. Estimating disease losses to the Australian wheat industry. Australasian Plant Pathology 38. https://doi.org/10.1071/AP09053

Nosanchuk, J.D., Casadevall, A., 2006. Impact of Melanin on Microbial Virulence and Clinical Resistance to Antimicrobial Compounds. Antimicrobial Agents and Chemotherapy 50, 3519–3528. https://doi.org/10.1128/aac.00545-06

Nosanchuk, J.D., Casadevall, A., 2003. The contribution of melanin to microbial pathogenesis. Cellular Microbiology 5, 203–223. https://doi.org/10.1046/j.1462-5814.2003.00268.x

Oliver, R., Friesen, T., Faris, J., Solomon, P., 2012. *Stagonospora nodorum:* From Pathology to Genomics and Host Resistance. Annual review of phytopathology 50, 23–43. https://doi.org/10.1146/annurev-phyto-081211-173019

Paolo, W.F., Dadachova, E., Mandal, P., Casadevall, A., Szaniszlo, P.J., Nosanchuk, J.D., 2006. Effects of disrupting the polyketide synthase gene WdPKS1 in *Wangiella* [Exophiala] *dermatitidis* on melanin production and resistance to killing by antifungal compounds, enzymatic degradation, and extremes in temperature. BMC Microbiology 6, 55. https://doi.org/10.1186/1471-2180-6-55

Pereira, D.A., McDonald, B.A., Brunner, P.C., 2017. Mutations in the *CYP51* gene reduce DMI sensitivity in *Parastagonospora nodorum* populations in Europe and China. Pest Management Science 73. https://doi.org/10.1002/ps.4486

Pereira, D.A.S., Ceresini, P., Castroagudín, V.L., Molina, L.M.R., Mesa, E.C., Negrisoli, M.M., Campos, S.N., Pegolo, M.E. de S., Takada, H.M., 2016. Population genetic structure of *Rhizoctonia oryzae-sativae* from rice in Latin America and its adaptive potential to emerge as pathogen on Urochloa pastures. Phytopathology 107, 121–131. https://doi.org/10.1094/phyto-05-16-0219-r

Pfeifer, B., Wittelsbürger, U., Ramos-Onsins, S.E., Lercher, M.J., 2014. PopGenome: An Efficient Swiss Army Knife for Population Genomic Analyses in R. Molecular Biology and Evolution 31, 1929–1936. https://doi.org/10.1093/molbev/msu136

Price, C.L., Parker, J.E., Warrilow, A.G., Kelly, D.E., Kelly, S.L., 2015. Azole fungicides - understanding resistance mechanisms in agricultural fungal pathogens. Pest Management Science 71, 1054–1058. https://doi.org/10.1002/ps.4029

Qin, C.-F., He, M.-H., Chen, F.-P., Zhu, W., Yang, L.-N., Wu, E.-J., Guo, Z.-L., Shang, L. P., Zhan, J., 2016. Comparative analyses of fungicide sensitivity and SSR marker variations indicate a low risk of developing azoxystrobin resistance in *Phytophthora infestans*. Scientific Reports 6, 20483. https://doi.org/10.1038/srep20483

Quaedvlieg, W., Verkley, G.J.M., Shin, H.-D., Barreto, R.W., Alfenas, A.C., Swart, W.J., Groenewald, J.Z., Crous, P.W., 2013. Sizing up Septoria. Studies in Mycology 75, 307–390. https://doi.org/10.3114/sim0017

R Core Team, 2019. R: A Language and Environment for Statistical Computing. R Foundation for Statistical Computing, Vienna, Austria.

Rehnstrom, A.L., Free, S.J., 1996. The isolation and characterization of melanin-deficient mutants of *Monilinia fructicola*. Physiological and Molecular Plant Pathology 49, 321–330. https://doi.org/10.1006/pmpp.1996.0057

Richards, J.K., Wyatt, N.A., Liu, Z., Faris, J.D., Friesen, T.L., 2017. Reference Quality Genome Assemblies of Three *Parastagonospora nodorum* Isolates Differing in Virulence on Wheat. G3: Genes, Genomes, Genetics 8, g3.300462.2017. https://doi.org/10.1534/g3.117.300462

Sanjak, J.S., Sidorenko, J., Robinson, M.R., Thornton, K.R., Visscher, P.M., 2018. Evidence of directional and stabilizing selection in contemporary humans. Proceedings of the National Academy of Sciences 115, 151–156. https://doi.org/10.1073/pnas.1707227114

Savary, S., Willocquet, L., Pethybridge, S., Esker, P., McRoberts, N., Nelson, A., 2019. The global burden of pathogens and pests on major food crops. Nature Ecology & Evolution 3, 430–439. https://doi.org/10.1038/s41559-018-0793-y

Shaw, M., Bearchell, S., Fitt, B., Fraaije, B., 2008. Long-term relationships between environment and abundance in wheat of *Phaeosphaeria nodorum* and *Mycosphaerella graminicola*. New Phytologist 177, 229–238. https://doi.org/10.1111/j.1469-8137.2007.02236.x

Singaravelan, N., Grishkan, I., Beharav, A., Wakamatsu, K., Ito, S., Nevo, E., 2008. Adaptive Melanin Response of the Soil Fungus *Aspergillus niger* to UV Radiation Stress at “Evolution Canyon”, Mount Carmel, Israel. PLoS ONE 3, e2993. https://doi.org/10.1371/journal.pone.0002993

Sommerhalder, R.J., McDonald, B.A., Zhan, J., 2006. The Frequencies and Spatial Distribution of Mating Types in *Stagonospora nodorum* Are Consistent with Recurring Sexual Reproduction. Phytopathology 96, 234–239. https://doi.org/10.1094/PHYTO-96-0234

Spitze, K., 1993. Population structure in *Daphnia obtusa:* quantitative genetic and allozymic variation. Genetics 135, 367–74.

Stefansson, T., Willi, Y., Croll, D., McDonald, 2014. An assay for quantitative virulence in *Rhynchosporium commune* reveals an association between effector genotype and virulence. Plant Pathology 63, 405–414. https://doi.org/10.1111/ppa.12111

Stefansson, T.S., McDonald, B.A., Willi, Y., 2013. Local adaptation and evolutionary potential along a temperature gradient in the fungal pathogen *Rhynchosporium commune*. Evolutionary Applications 6, 524–534. https://doi.org/10.1111/eva.12039

Stukenbrock, E.H., Banke, S., McDonald, B.A., 2006. Global migration patterns in the fungal wheat pathogen *Phaeosphaeria nodorum*. Molecular Ecology 15, 2895–2904. https://doi.org/10.1111/j.1365-294x.2006.02986.x

Stukenbrock, E.H., McDonald, B.A., 2008. The Origins of Plant Pathogens in AgroEcosystems. Annual Review of Phytopathology 46, 75–100. https://doi.org/10.1146/annurev.phyto.010708.154114

Sturm, R.A., Duffy, D.L., 2012. Human pigmentation genes under environmental selection. Genome Biology 13, 248. https://doi.org/10.1186/gb-2012-13-9-248

Tonsor, S.J., Elnaccash, T.W., Scheiner, S.M., 2013. Developmental instability is genetically correlated with phenotypic plasticity, constraining heritability, and fitness. Evolution 67, 2923–2935. https://doi.org/10.1111/evo.12175

Wheeler, M.H., Bell, A.A., 1988. Melanins and Their Importance in Pathogenic Fungi, in: Springer New York. pp. 338–387. https://doi.org/10.1007/978-1-4612-3730-3_10

Whitlock, M.C., 2008. Evolutionary inference from QST. Molecular Ecology 17, 1885–1896. https://doi.org/10.1111/j.1365-294X.2008.03712.x

Wickham, H., 2009. Ggplot2: Elegant Graphics for Data Analysis, 2nd ed. Springer Publishing Company, Incorporated.

Willi, Y., Frank, A., Heinzelmann, R., Kälin, A., Spalinger, L., Ceresini, P.C., 2011. The adaptive potential of a plant pathogenic fungus, *Rhizoctonia solani* AG-3, under heat and fungicide stress. Genetica 139, 903. https://doi.org/10.1007/s10709-011-9594-9

Yampolsky, L., Schaer, T., Ebert, D., 2013. Adaptive phenotypic plasticity and local adaptation for temperature tolerance in freshwater zooplankton. Proceedings of the Royal Society B: Biological Sciences 281, 20132744–20132744. https://doi.org/10.1098/rspb.2013.2744

Yang, L., Zhu, W., Wu, E., Yang, C., Thrall, P.H., Burdon, J.J., Jin, L., Shang, L., Zhan, J., 2016. Trade-offs and evolution of thermal adaptation in the Irish potato famine pathogen *Phytophthora infestans*. Molecular Ecology 25, 4047–4058. https://doi.org/10.1111/mec.13727

Zearfoss, A., Cowger, C., Ojiambo, P., 2011. A Degree-Day Model for the Latent Period of Stagonospora nodorum Blotch in Winter Wheat. Plant Disease 95, 561–567. https://doi.org/10.1094/pdis-09-10-0651

Zhan, J., Linde, C.C., Jürgens, T., Merz, U., Steinebrunner, F., McDonald, B.A., 2005. Variation for neutral markers is correlated with variation for quantitative traits in the plant pathogenic fungus *Mycosphaerella graminicola*. Molecular Ecology 14, 2683–2693. https://doi.org/10.1111/j.1365-294x.2005.02638.x

Zhan, J., McDonald, B., 2011. Thermal adaptation in the fungal pathogen *Mycosphaerella graminicola*. Molecular Ecology 20. https://doi.org/10.1111/j.1365-294X.2011.05023.x

Zhu, W., Zhan, J., McDonald, B.A., 2018. Evidence for local adaptation and pleiotropic effects associated with melanization in a plant pathogenic fungus. Fungal Genetics and Biology. https://doi.org/10.1016/j.fgb.2018.04.002

